# Merger of *Betula tatewakiana* (Betulaceae) from northern Japan to northeast Asian *B. ovalifolia* based on ploidy level

**DOI:** 10.1101/857359

**Authors:** Yuki Shiotani, Tomoko Fukuda, Elena A. Pimenova, Ekaterina A. Petrunenko, Pavel V. Krestov, Svetlana N. Bondarchuk, Yoko Nishikawa, Takashi Shimamura, Yoshiyasu Fujimura, Koh Nakamura

## Abstract

It has been controversial whether *Betula tatewakiana*, a dwarf birch distributed in Hokkaido of northern Japan, is an endemic species or a synonym of *B. ovalifolia* broadly distributed in northeast Asia. The endemic hypothesis is based on the idea that *B. tatewakiana* is diploid while *B. ovalifolia* is tetraploid and that they are separated based on the ploidy level; however no chromosome data have actually been published before. Resolving the taxonomic problem is crucial also in judging the conservation priority of *B. tatewakiana* in a global perspective. Our chromosome observation revealed that *B. tatewakiana* is tetraploid as well as *B. ovalifolia*. Collaterally, we conducted morphological observation and clarified that *B. tatewakiana* is morphologically identical to *B. ovalifolia* in white hairs and dense resinous glands respectively on adaxial and abaxial leaf surfaces, based on which they are different from closely related species in the same section *Fruticosae*. We concluded that the hypothesis that *B. tatewakiana* is a Hokkaido endemic based on the ploidy level is not supported and that *B. tatewakiana* should be merged with *B. ovalifolia.*

## Introduction

*Betula ovalifolia* Rupr. (1857: 378) is a dwarf birch found in wetlands (Gray 1996, Li & Skvortsov 1999). It grows only up to ca. 2 m tall (Li & Skvortsov 1999) and reproduces not only by seeds but also asexually by branching near ground level (Tabata 1966). This species is widely distributed in northeast Asia, i.e., Russian Far East (southern Khabarovsky Krai, Primorsky Krai, Amur Oblast, Jewish Autonomous Oblast), northeast China (Heilongjiang, Changbai Shan, Nei Mongol), north Korea, and northern Japan (Hokkaido) (Fig. 1; Gray 1996, Li & Skvortsov 1999). However, taxonomic treatment of *B. ovalifolia* from Japan, and thereby its occurrence in Japan, has been controversial. In the first report from Japan, it was treated as an endemic species of Hokkaido and named *B. tatewakiana* M.Ohki & S.Watan. (1959: 9). Afterward, it had been treated as a variety of *B. humilis* Schrank (1789: 420) (Murata 1978). Soon after that, Hara (1979) claimed that this species is identical to *B. ovalifolia*, but no data were presented. Since then, although this opinion is widely accepted in pictorial books and floras in Japan (Murata 1979, Ito 1981, Ohba 2006, Takahashi 2015), the idea to support *B. tatewakiana* was claimed again (Watanabe 1995) and the taxonomic problem still remains (Takahashi 2013, Takahashi et al. 2013, Nemoto 2016). Here, we tentatively used the name *B. tatewakiana* and later discuss its taxonomy and proper name based on our result.

**FIGURE 1.**
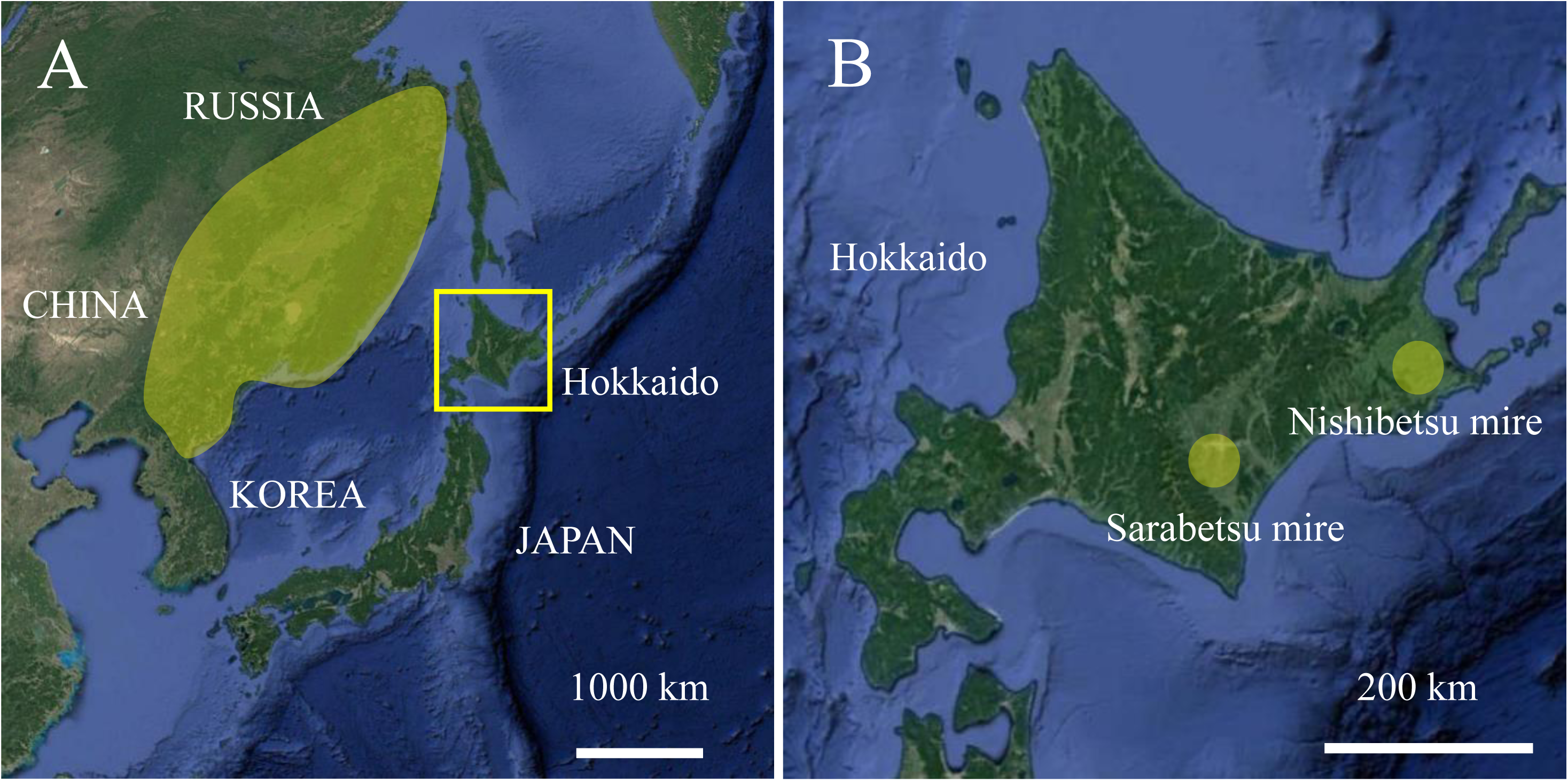
Species distribution ranges of *Betula ovalifolia* (A) and *B. tatewakiana* (B).

The taxonomic problem of *B. tatewakiana* and *B. ovalifolia* stems from the confusion in their ploidy level. Watanabe (1995) claimed that *B. tatewakiana* is diploid while *B. ovalifolia* is tetraploid and he recognized *B. tatewakiana* as the Japanese endemic restricted to Hokkaido. This idea, however, was originally reported in a conference abstract without images of the chromosomes (Watanabe & Somego 1991) and has never been published, but repeatedly mentioned in following studies (Nemoto 2016). On the other hand, Nagamitsu (2004) did not separate the two species and treated *B. ovalifolia* from Hokkaido as a tetraploid species based on Provatoba (1995), which actually didn’t report the chromosome number of *B. ovalifolia* but of a hybrid between *B. ovalifolia* and *B. exilis*. A flow cytometric study of the genome size evolution in the genus *Betula* suggested that *B. ovalifolia* from the Asian continent is tetraploid (Wang *et al*. 2016), but no information based on chromosome observation exist about the ploidy level of *B. tatewakiana* and *B. ovalifolia*.

In this study, to resolve the taxonomic problem of *B. tatewakiana*, we conducted chromosome observation and determined the ploidy level. We collaterally conducted morphological observation of *B. tatewakiana*. Regarding *B. ovalifolia*, there are two closely related species in the same section *Fruticosae*, i.e., *B. humilis* Schrank (1789: 420) and *B. fruticosa* Pall. (1776: 758) and *B. ovalifolia* is distinguished from the two species by white hairs on adaxial leaf surface (vs. glabrous in *B. humilis* and *B. fruticosa*) and by densely resinous glands on abaxial leaf surface (vs. lack of glands in *B. humilis*) (Kuzeneva 1985, Li & Skvortsov 1999). In previous studies which did not accept *B. tatewakiana*, these traits have not been well compared between *B. tatewakiana* and *B. ovalifolia*. Resolving the taxonomic problem and assess the endemic status of *B. tatewakiana* would also help planning its conservation. *Betula tatewakiana* is distributed only in two localities in Japan, i.e., Sarabetsu and Nishibetsu mires in eastern Hokkaido (Fig. 1-B). As a result of the exploitation of the mires, remaining habitats are only 3 and 16 ha in Sarabetsu and Nishibetsu mires, respectively (Takahashi 2013). Open ditches excavated inside and outside the mires are increasingly drying the habitats of *B. tatewakiana* and thereby it is red-listed at national and prefectural levels (Hokkaido 2001, Ministry of the Environment of Japan 2018). Whether it is endemic or not is related to its conservation priority in a global perspective; on the other hand, if it is the same species as *B. ovalifolia* broadly distributed in northeast Asia, effective conservation should be planned considering genetic connectivity with conspecific populations abroad. This study is expected to provide basic information essential for the conservation of the species.

## Material & Methods

### Determination of ploidy level

We collected seeds of *B. tatewakiana* from six and five individuals from Sarabetsu and Nishibetsu mires in Hokkaido, Japan; seeds of *B. ovalifolia* were collected from one individual in Sikhote-Alin Nature Reserve in Primorsky Krai, Russian Far East (Table 1). Collected seeds were dried with silica gel and stored at 4°C. Seeds were sowed on vermiculite and germinated at 25°C day / 8°C night condition for two weeks. After germination, root tips were collected and pretreated by 0.002 M 8-hydroxyquinoline solution for 24 hours at 4°C in dark condition. Next, the root tips were fixed by Farmer’s solution (glacial acetic acid : 99 % ethanol = 1 : 3) at 4°C in dark condition. After fixation, the root tips were macerated in 1 N HCl for 18 minutes and stained with 1 % aceto-orcein for 5 minutes and squashed on a slide. Metaphase chromosomes were observed using an optical microscope Zeiss Axio Imager A1 (Carl Zeiss, Jena, Germany), and pictured by Anyty 3R-DKMC01 (3R solution corp., Fukuda, Japan).

**TABLE 1.** Chromosome counts of *Betula tatewakiana* and *B. ovalifolia*.

| **Taxon** | **Sampling site** | **Chromosome counts** | **Voucher no.** |
| --- | --- | --- | --- |
| *B. tatewakiana* | Sarabetsu mire, Sarabetsu village,<br>Hokkaido, Japan | 56 | HUBG* 14746 A, E, H |
|  |  | 52 | HUBG 14746 B, D |
|  |  | 50 | HUBG 14746 F |
|  | Nishibetsu mire, Betsukai town,<br>Hokkaido, Japan | 56 | Yuki Shiotani 1, 26, 29, 30 (SAPT) |
|  |  | 53 | Yuki Shiotani 27 (SAPT) |
| *B. ovalifolia* | Sikhote-Alin Nature Reserve,<br>Terney, Primorsky Krai, Russia | 56 | Koh Nakamura 14198 (SAPT) |

\*HUBG: living collections of Hokkaido University Botanic Garden

### Morphological observations

To elucidate whether *B. tatewakiana* is morphologically identical to *B. ovalifolia* or not, we observed the key traits in the section *Fruticosae*: white hairs and dense resinous glands respectively on adaxial and abaxial leaf surfaces. For *B. tatewakiana*, specimens examined were the holotype of *B. tatewakiana* (H. Suzuki & M. Ohki, s.n. with handwriting “Type”) in the herbarium of Hokkaido University Museum (SAPS) and our collections of 51 and 45 plants from Sarabetsu and Nishibetsu mires, that were deposited in the herbarium of Hokkaido University Botanic Garden (SAPT) (Appendix 1). For *B. ovalifolia*, our collections of 38 specimens from Primorsky Krai in Russia Far East were used (SAPT, Appendix 1).

## Results

### Ploidy level

Somatic chromosomes at metaphase were approximately 1.0 µm long in both *B. tatewakiana* (Fig. 2 A–D) and *B. ovalifolia* (Fig. 2 E, F). The centromere positions could not be determined because of the small sizes of the chromosomes. The result of chromosome counts is summarized in Table 1. In *B. tatewakiana* from Sarabetsu mire, 3 individuals had 56 chromosomes (HUBG 14746 A, 14746 E, and 14746 H), 2 individuals had ca. 52 chromosomes (HUBG 14746 B, D) and 1 individual had ca. 50 chromosomes (HUBG 14746 F). In *B. tatewakiana* from Nishibetsu mire, 4 individuals had 56 chromosomes (Yuki Shiotani 1, 26, 29, 30) and 1 individual had ca. 53 chromosomes (Yuki Shiotani 27). In *B. ovalifolia* from Primorsky Krai, 1 individual had 56 chromosomes (Koh Nakamura 14198).

**FIGURE 2.**
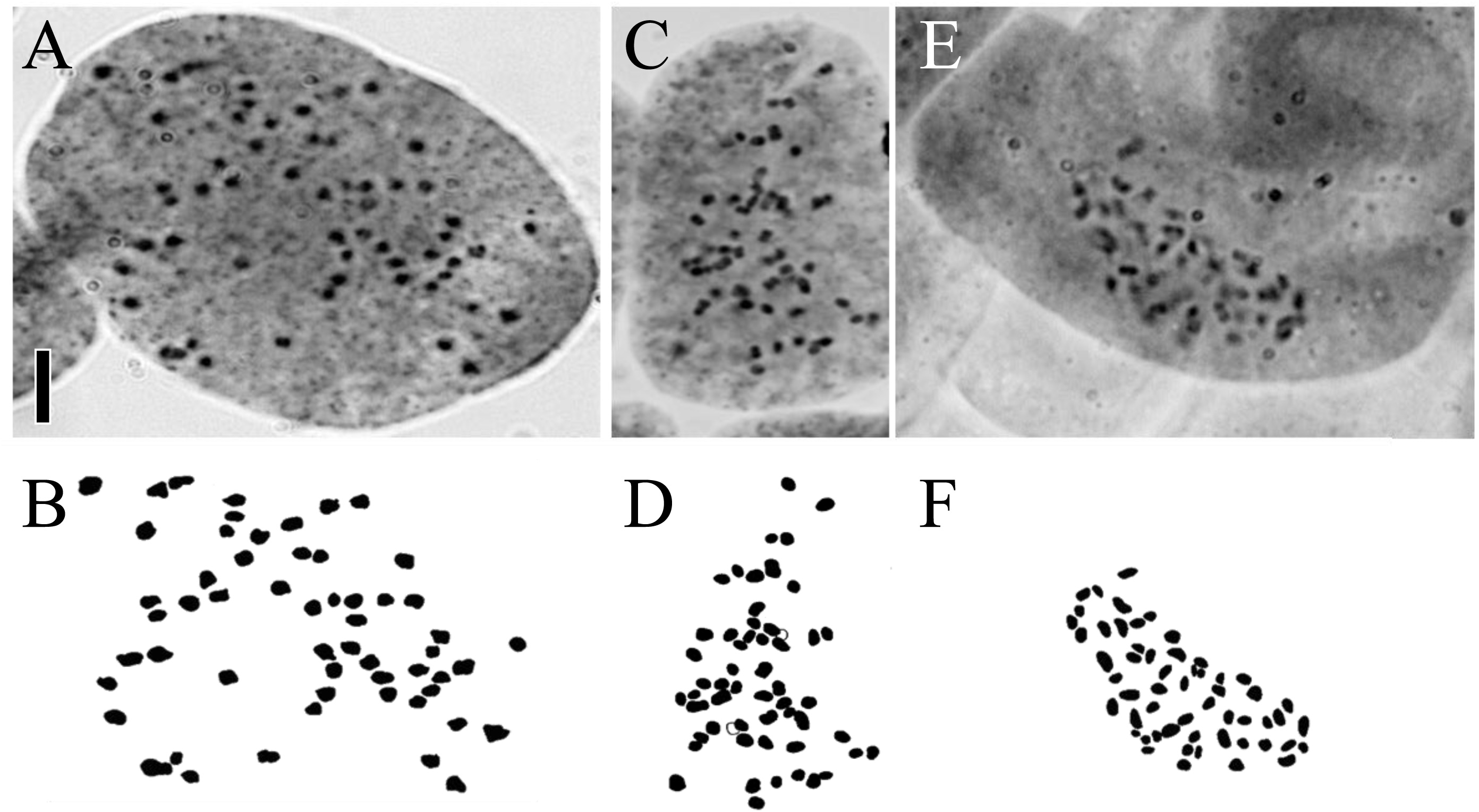
Somatic chromosomes at metaphase of *B. tatewakiana* and *B. ovalifolia*. Photomicrographs of *B. tatewakiana* from Sarabetsu mire (A, 2*n* = 56: HUBG 14746 A and Nishibetsu mire (C, 2*n* = 56: Yuki Shiotani 29), and *B. ovalifolia* from Primorsky Krai (E, 2*n* = 56: Koh Nakamura 14198) are shown. B, D, and F are drawings of A, C, and E, respectively. Scale bar is 5 µm.

### Morphological traits

The holotype of *B. tatewakiana* had white hairs and dense resinous glands respectively on adaxial and abaxial leaf surface (Fig. 3 A, B). Our collections of *B. tatewakiana* also had white hairs and dense resinous glands on adaxial and abaxial leaf surface, respectively (Fig. 3 C, D) and no morphological difference was recognized between the samples from Sarabetsu and Nishibetsu mires. In *B. ovalifolia*, our collections from Primorsky Krai had white hairs and dense resinous glands respectively on adaxial and abaxial leaf surface as well as *B. tatewakiana* (Fig. 3 E, F).

**FIGURE 3.**
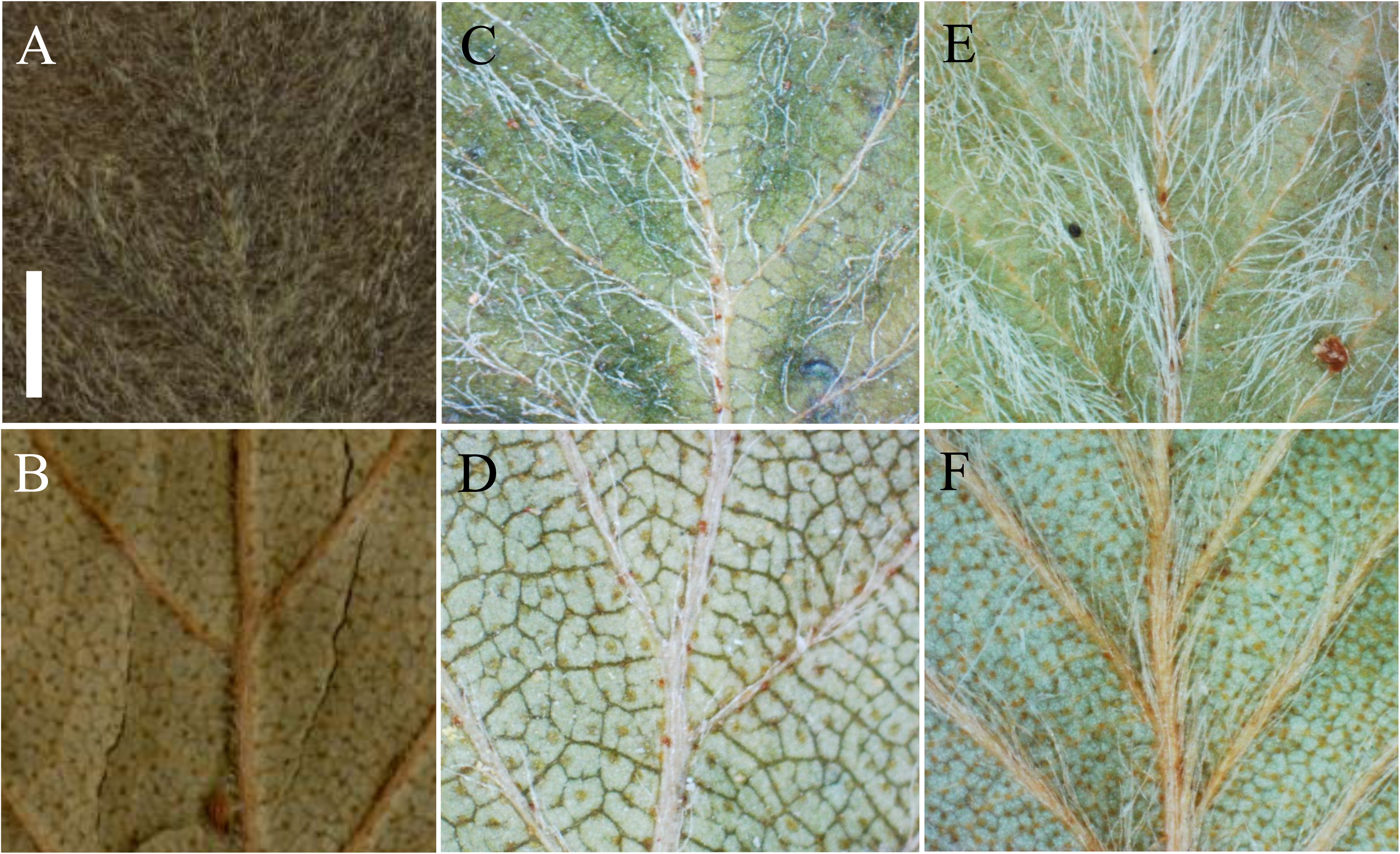
Leaf traits of *B. tatewakiana* and *B. ovalifolia*. White hairs on adaxial leaf surface (A, C, E) and densely resinous glands on abaxial leaf surface (B, D, F) are shown for the holotype of *B. tatewakiana* (H. Suzuki & M. Ohki, s.n., A & B), *B. tatewakiana* of our collection (Yuki Shiotani 38, C & D), and *B. ovalifolia* in Russia (Koh Nakamura 14188, E & F). Scale bar is 1 mm.

## Discussion

### Merger of B. tatewakiana to B. ovalifolia

In our chromosome observation, the samples of *B. tatewakiana* from Sarabetsu and Nishibetsu mires had 2*n* = 50 (one sample), 52 (two samples), 53 (one sample), and 56 (seven samples) chromosomes (Table 1). The chromosomes were too small (approximately 1.0 µm long) to observe clearly and the chromosome count variation may need further verification; however, it would be safe to say that *B. tatewakiana* is tetraploid because the basic chromosome number is 14 in the genus *Betula* (Erikkson & Jonsson 1986) and the diploid count should be 2*n* = 28. Watanabe & Somego (1991) reported that *B. tatewakiana* is diploid, although no images of the chromosomes were presented. Thus, the possibility that there are both diploid and tetraploid in *B. tatewakiana* is not totally denied. However, his report was gametophytic count and according to the author Watanabe the chromosome image was unclear (personal communication). For this reason, *B. tatewakiana* is highly likely to be tetraploid. Our chromosome count of *B. ovalifolia* was 2*n* = 56. This is consistent with the flow cytometric study that suggested that *B. ovalifolia* from the Asian continent is tetraploid (Wang *et al*. 2016). Therefore, the idea to separate *B. tatewakiana* from *B. ovalifolia* based on the ploidy level (Watanabe & Somego 1991, Watanabe 1995) is not supported because both species are tetraploid. Hence, *B. tatewakiana* should be merged to *B. ovalifolia*. The observation of the morphological traits also supports the merger of *B. tatewakiana* to *B. ovalifolia*. The two species are morphologically identical in white hairs and dense resinous glands respectively on adaxial and abaxial leaf surfaces, based on which they are different from closely related dwarf birch species in the same section *Fruticosae*.

### Implications for conservation

*Betula tatewakiana* is recognized as a synonym of *B. ovalifolia* as discussed above, and thereby it is not a Japanese endemic species. Hereafter the Hokkaido populations are called *B. ovalifolia*. Because *B. ovalifolia* is broadly distributed in northeast Asia, i.e., Russian Far East, northeast China, north Korea, and northern Japan, the conservation priority of the species may not be high in a global perspective. On the other hand, the Hokkaido populations represent only island populations disjunct from continental populations. The species had likely moved southward during glacial periods and retreated northward in warmer periods, and the Hokkaido populations are considered to be relict populations (Takahashi 2013). The Hokkaido populations can be reproductively isolated from the continental populations and can have a unique gene pool that deserves conservation. Also, domestically in Japan, *B. ovalifolia* is distributed only in Sarabetsu and Nishibetsu mires in Japan and deserves conservation as national resource. If there exists geneflow among Hokkaido and continental populations, effective conservation should be planned considering genetic connectivity with populations abroad. Population genetics of *B. ovalifolia* in northeast Asia for conservation is the topic of our future investigation.

## Acknowledgement

We thank staff of Hokkaido prefecture, Betsukai town, and Sarabetsu village for their helps in sampling permission. This study was supported in part by Grants-in-Aid for Scientific Research, KAKENHI to K.N. (16K18596) and a research grant from the Mitsui & Co. Environment Fund to K.N. (R15-0067).

## Appendix1 Specimens for morphological observation

***Betula tatewakiana***

Japan, Hokkaido: Sarabetsu village, 18 August 1958, (H. Suzuki & M. Ohki, s.n. with handwriting “Type”, SAPS); Sarabetsu mire, 5 June 2017, Yuki Shiotani 65–115 (51 specimens, SAPT); Nishibetsu mire, 7 June 2017, Yuki Shiotani 1–26, 31–49 (45 specimens, SAPT)

***B. ovalifolia***

Russia, Primorsky Krai: Terney, 22 July 2016, Koh Nakamura 14169–14195, 14197, 14198 (29 specimens, SAPT); Terney, 23 July 2016, Koh Nakamura 14289–14297 (9 specimens, SAPT)

## References

Erikkson, G. & Jonsson, A. (1986) A review of the genetics of *Betula*. Scandinavian Journal of Forest Research 1: 421–434.

Gray, S.F. (1996) Betula. In: Charkevicz, S.S. (Ed.) Plantae Vasculare Orientis Extremi Sovietici 8. NAUK, Moscow, pp. 13–24.

Hara, H. (1979) Comments on the East Asiatic plants 6. Journal of Japanese Botany 54: 1–9.

Hokkaido (2001) Hokkaido red list (plants). Hokkaido Government, Sapporo. Available from: http://www.pref.hokkaido.lg.jp/ks/skn/grp/03/redlist1.pdf (accessed: 8 October 2019).

Ito, K. (1981) Alpine Plants in Hokkaido. Seibundo Shinkosha, Tokyo, pp. 230.

Kuzeneva, O.I. (1985) Betula. In: Komarov, V.L. (Ed.) Flora of the U.S.S.R. 5. NAUK, Moscow. Available from: https://archive.org/stream/floraofussr05bota#page/n5/mode/2up (accessed: 8 October 2019).

Li, P. & Skvortsov, A.K. (1999) Betulaceae. In: Wu, Z.Y. & Raven, P.H. (Eds.) Flora of China 4. Science Press, Beijing, and Missouri Botanical Garden Press, St. Louis. pp. 286–313.

Maack, R & Ruprecht, J.F. (1857) Premières nouvelles botaniques des rives de l’Amour. In: Bulletin de la Classe Physico-Mathématique 15. l’Académie Impériale des Sciences, St. Peterburg, pp. 378.

Ministry of the Environment, Government of Japan (2018) Ministry of the Environment red list 2017. Ministry of the Environment, Government of Japan, Tokyo. Available from: https://www.env.go.jp/press/files/jp/109278.pdf (accessed: 8 October 2019).

Murata, G. (1978) Taxonomical Notes 12. Acta Phytotaxonomica et Geobotanica 29: 95–105.

Murata, G. (1979) Betulaceae. In: Kitamura, S. & Murata, G. (Eds.) Colored illustrations of herbaceous plants of Japan (Choripetalae). Hoikusha, Osaka, pp. 284–304.

Nagamitsu, T., Kawahara, T. & Hotta, M. (2004) Phenotypic variation and leaf fluctuating asymmetry in isolated populations of an endangered dwarf birch *Betula ovalifolia* in Hokkaido, Japan. Plant Species Biology 19: 13–21.

Nemoto, T. (2016) Betulaceae. In: Ohashi, H., Kadota, Y., Murata, J., Yonekura, K. & Yahara, K. (Eds.) Wild Flowers of Japan II revised new edition. Heibonsha, Tokyo, pp. 107–119.

Ohba, H. (2006) Betula L. In: Iwatsuki, K., Boufford David, E. & Ohba, H. (Eds.) Flora of Japan. Angiospermae, Dicotyledoneae, Archichlamydeae (a) vol IIa. Koudansha, Tokyo, pp. 31–37.

Pallas, P.S. (1776) Reise durch Verschiedene Provinzen des Russischen Reichs 3. Academie der Wissenschaften, St. Peterburg. 758 pp.

Probatova, N.S. & Sokolovskaya, A.P. (1995) Chromosome numbers in some species of vascular plants from the Russian Far East. Botanicheskii Zhurnal 80: 85–88.

Schrank, F.P. (1789) Baiersche Flora. Johann Baptist Strobl, München, 420 pp.

Tabata, H. (1966) A contribution to the biology of Japanese birches. Memories of the College of Science, University of Kyoto, Series B, Biology 32: 239–271.

Takahashi, H. (2013) Report on the habitats of *Betula tatewakiana* in Betsukai-cho. In: Educational committee of Betsukai-cho (Ed.) *Report on the habitats of Betula tatewakiana in Nishibetsu mire, Natural monument of Hokkaido*. Educational committee of Betsukai-cho, Betsukai, pp. 13–15.

Takahashi, H. (2015) Betula ovalifolia. In: Yahara, T., Fujii, S., Ito, M. & Ebihara, A. (Eds.) Red data plants. Yama-kei Publishers, Tokyo, p. 212.

Takahashi, H., Oida, K., Takahashi, M. & Yonaha, T. (2013) *Betula tatewakiana* habitats in Betsukai-cho. In: Educational committee of Betsukai-cho (Ed.) Report on the habitat of Betula tatewakiana in Nishibetsu mire, Natural monument of Hokkaido. Educational committee of Betsukai-cho, Betsukai, pp. 5–12.

Wang, N., McAllister, H.A., Bartlett, P.R. & Buggs, R.J.A. (2016) Molecular phylogeny and genome size evolution of the genus *Betula* (Betulaceae). Annals of Botany 117: 1023–1035.

Watanabe, T. (1995) Betulaceae. In: Iwatsuki, K., Ohba, H., Shimizu, T., Hotta, M., Prance, T.G. & Raven, H.P. (Eds.) Asahi encyclopedia. The world of plants 8, The Asahi Shimbun Tokyo, pp. 98–119.

Watanabe, T. & Ohki, M. (1959) A new species of *Betula* from Hokkaido. Journal of Japanese Botany 34: 329–332.

Watanabe, T. & Somego, M. (1991) Chromosome numbers of two species of subgenus *Chamaebetula*, genus *Betula* from Japan. In: Botanical Society of Japan (Ed.) *Proceedings of the 56th annual meeting of the Botanical Society of Japan*. Botanical Society of Japan, Tokyo, p. 331.

